# Oxa1L buffers mitochondrial vulnerability by coupling translation to membrane insertion

**DOI:** 10.64898/2025.12.22.696118

**Authors:** Xurui Shen, Rui He, Huiwei Zhong, Peixin Sun, Jinlun Kylian Zhang, Yue Zhang, Hanting Yang

## Abstract

Mitochondrial respiratory chain integrity relies on the coordinated synthesis and membrane insertion of mitochondrially encoded proteins. How limitations in this process contribute to mitochondrial dysfunction under stress remains poorly understood. Here, we identify the mitochondrial insertase Oxa1L as a key buffering factor that modulates mitochondrial vulnerability under Parkinsonian stress conditions. Transcriptomic analyses and cellular stress models reveal pronounced disruption of oxidative phosphorylation pathways accompanied by reduced Oxa1L abundance. Functional gain- and loss-of-function analyses demonstrate that Oxa1L actively influences mitochondrial membrane potential, reactive oxygen species levels, and ATP production during stress. Notably, Oxa1L selectively governs the abundance of mitochondrially encoded respiratory chain subunits, indicating that co-translational membrane insertion represents a rate-limiting step in mitochondrial protein biogenesis. Structural analyses position Oxa1L at the mitoribosomal exit site, providing a spatial framework for coupling translation to inner membrane insertion. Together, our findings uncover a critical layer of mitochondrial quality control at the level of co-translational insertion and establish Oxa1L as a determinant of mitochondrial resilience under stress.

## Introduction

Misfolded proteins within mitochondria are increasingly recognized as important contributors to neurodegenerative diseases ^1–3^. In dopaminergic neurons of the substantia nigra, protein misfolding, abnormal aggregation, and fibrillation trigger a cascade of pathological events that disrupt mitochondrial protein transport ^3, 4^. Such disturbances compromise mitochondrial integrity and respiratory chain function, leading to impaired energy metabolism, oxidative phosphorylation (OXPHOS) dysfunction, and ultimately neuronal degeneration and death, thereby contributing to the onset of neurodegenerative diseases such as Parkinson’s disease (PD)^5^ ^6, 7^.

Within mitochondria, proper expression and membrane integration of respiratory chain components are essential for maintaining OXPHOS activity. Nascent peptide chains synthesized by mitochondrial ribosomes are inserted into the inner mitochondrial membrane through a co-translational translocation mechanism, which ensures the correct localization and assembly of mitochondrial genome-encoded proteins^8–10^. This tightly coupled process minimizes exposure of hydrophobic polypeptides to the aqueous matrix and is therefore critical for preserving mitochondrial proteostasis and bioenergetic function.

The mitochondrial inner membrane insertase Oxa1L (Oxidase assembly 1-like) plays a central role in this process. Oxa1L localizes to the inner mitochondrial membrane and facilitates the co-translational insertion of nascent polypeptide chains encoded by the mitochondrial genome by directly interacting with mitochondrial ribosomes^9, 11–15^.

In addition to its role in inserting mitochondrial genome–encoded proteins, Oxa1 is also involved in the inner membrane sorting of a subset of nuclear-encoded proteins. Many nuclear-encoded inner membrane precursors are first imported into the mitochondrial matrix and processed peptidases, after which they are either translocated into the matrix via the presequence-associated motor (PAM) or inserted into the inner membrane through Oxa1-mediated pathways^16–18^. Although structures of the Oxa1L-mitoribosome complexes have been reported, their resolution remains limited, and the molecular determinants governing Oxa1L–ribosome coupling during co-translational membrane insertion are still incompletely understood^19–22^. Moreover, how this insertion machinery impacts mitochondrial function under pathological stress conditions remains largely unexplored.

Genetic studies have revealed that mutations in Oxa1L cause mitochondrial dysfunction and human neurological disorders^12, 23–26^. In affected patients, Oxa1L expression is reduced in brain and skeletal muscle tissues, accompanied by decreased abundance or activity of mitochondrial OXPHOS complexes IV and V, leading to neurodevelopmental delay and motor impairment^23^. These observations underscore the physiological importance of Oxa1L-mediated protein insertion for mitochondrial function. However, whether dysregulation of Oxa1L contributes to mitochondrial dysfunction in PD has not been investigated.

In this study, we show that MPP^+^-induced Parkinsonian stress leads to marked downregulation of Oxa1L, resulting in reduced expression of mitochondrial genome–encoded OXPHOS subunits, impaired oxidative phosphorylation, and disrupted energy metabolism. Through combined loss- and gain-of-function analyses, structural characterization of the Oxa1L–mitoribosome complex, and systematic truncation studies, we identify critical regions of Oxa1L contribute to co-translational membrane insertion and mitochondrial protection. Our findings reveal Oxa1L-dependent protein insertion as a previously underappreciated determinant of mitochondrial functional homeostasis and provide new mechanistic insight into mitochondrial dysfunction in PD pathogenesis.

## Results

### Altered Oxa1L expression in an MPP^+^-induced Parkinsonian cell model

To examine mitochondrial responses to Parkinsonian stress, SH-SY5Y cells were treated with MPP^+^ and subjected to RNA sequencing analysis. Differential expression profiling revealed widespread transcriptional alterations, among which approximately 6.5% of the differentially expressed genes were mitochondria-related (Fig. 1A, B) , indicating a pronounced impact of MPP^+^ treatment on mitochondrial pathways. Consistent with this observation, KEGG pathway enrichment analysis identified oxidative phosphorylation (OXPHOS) as one of the most significantly affected pathways (Fig. 1C). Gene Ontology (GO) enrichment analysis further revealed marked dysregulation of mitochondrial inner membrane–associated processes, including mitochondrial translation, inner mitochondrial protein complex organisation, and mitochondrial respiratory chain complex assembly (Fig. 1D).

**Figure 1.**
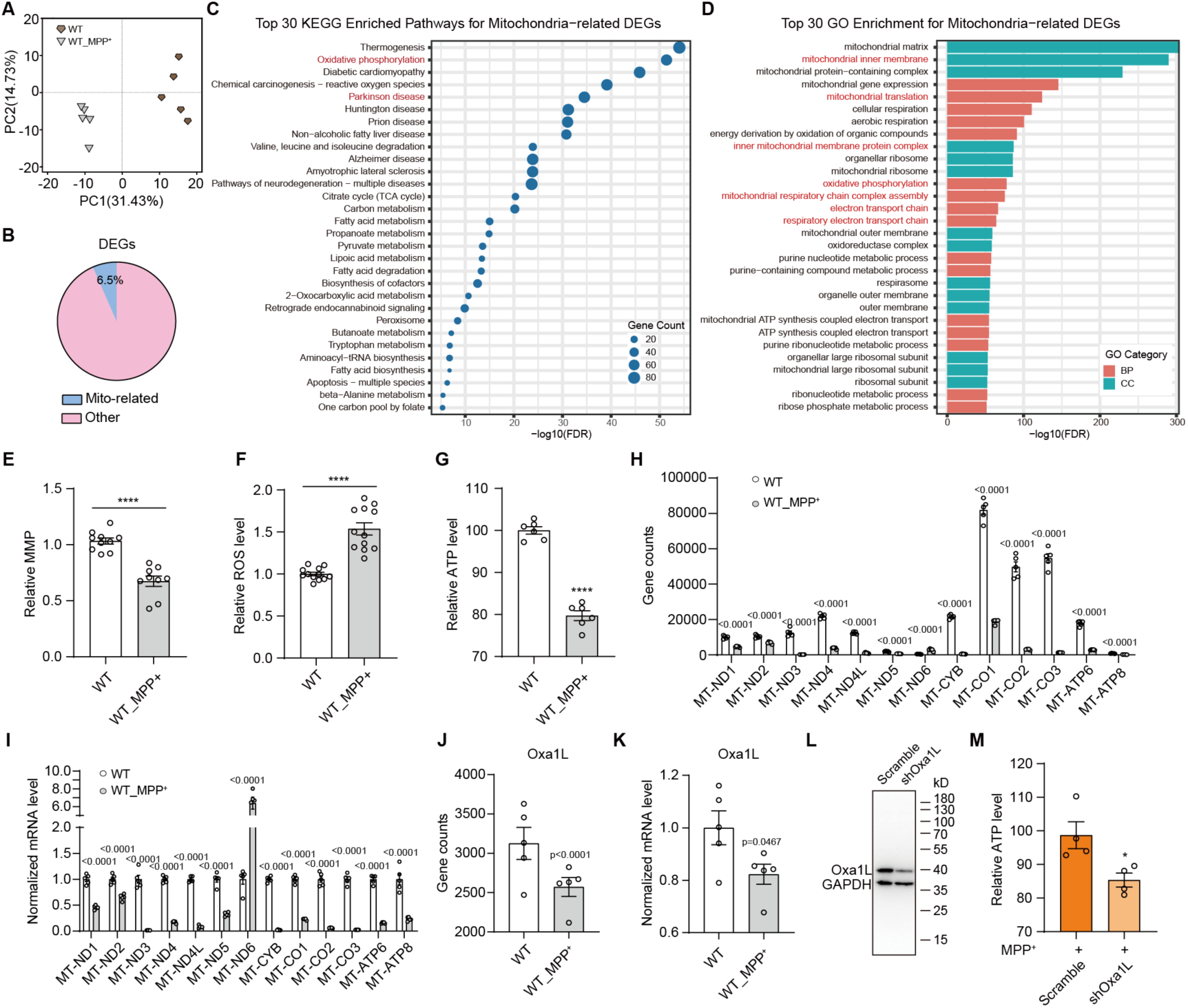
Mitochondrial dysfunction and altered Oxa1L expression in an MPP^+^-induced stress model. **A** Principal component analysis (PCA) of RNA-seq data from control (WT) and MPP^+^-treated (WT_MPP^+^) SH-SY5Y cells. **B** Proportion of mitochondria-related genes among all differentially expressed genes (DEGs). **C** KEGG pathway enrichment of mitochondria-related DEGs, highlighting oxidative phosphorylation (OXPHOS). **D** Gene Ontology enrichment showing dysregulation of mitochondrial–associated processes. **E-G** Mitochondrial membrane potential (E), intracellular ROS levels (F), and ATP levels (G) in control and MPP^+^-treated cells. **H, I** Gene counts (H) and normalized mRNA levels (I) of the 13 mitochondrial genome-encoded genes. **J, K** RNA-seq–based gene counts (**J**) and normalized mRNA levels (**K**) of Oxa1L. **L** Immunoblotting analysis of Oxa1L expression. **M** ATP levels in scramble and shOxa1L cells following MPP^+^ treatment. (Data represent mean ± SEM; statistical significance as indicated, *\*\*\*\*p < 0.0001*; *\*\*\*p < 0.001*; *\*\*p < 0.01*; *\*p < 0.05*; ns not significant).

We next assessed whether these transcriptional changes were reflected at the functional level of mitochondria. MPP^+^ exposure led to a reduction in mitochondrial membrane potential (MMP), together with elevated intracellular reactive oxygen species (ROS) levels and pronounced decrease in cellular ATP content (Fig. 1E-G), indicating compromised mitochondrial OXPHOS activity.

Because proper assembly of OXPHOS complexes requires coordinated expression of both nuclear- and mitochondrial genome-encoded subunits^27, 28^, we examined transcription of the 13 mitochondrial genome-encoded genes. With the exception of MT-ND6, which displayed modest upregulation, all other mitochondrial transcripts were reduced following MPP^+^ treatment (Fig. 1H, I), suggesting a global suppression of mitochondrial genome transcription under Parkinsonian stress.

Oxa1L is a mitochondrial inner membrane insertase contribute to the co-translational insertion of mitochondrially encoded proteins^11^. In line with the observed defects in mitochondrial gene expression and function, Oxa1L mRNA levels were decreased in MPP^+^-treated cells (Fig. 1J, K). Moreover, knockdown of Oxa1L further reduced intracellular ATP levels in the presence of MPP^+^ (Fig. 1L, M). Collectively, these data indicate that Oxa1L expression is downregulated in an MPP^+^-induced PD cell model and point to a potential contribution of Oxa1L loss to mitochondrial dysfunction.

### Consequences of Oxa1L deficiency for mitochondrial function under Parkinsonian stress

To assess whether Oxa1L modulates mitochondrial vulnerability to MPP^+^, we generated stable SH-SY5Y cell lines expressing shRNAs targeting Oxa1L (shOxa1L) (Fig. 2A, B). Comparative transcriptomic analysis between shOxa1L and scramble control cells revealed extensive gene expression differences following MPP^+^ treatment. GO enrichment analysis showed that genes reduced upon Oxa1L knockdown were preferentially associated with mitochondrial processes, including oxidative stress responses, protein targeting to mitochondria, and inner mitochondrial membrane organization (Fig. 2C, highlighted in red). Pathway-level analysis further supported these findings, as KEGG enrichment revealed deeper suppression of the oxidative phosphorylation pathway in Oxa1L-deficient cells relative to controls under MPP^+^ exposure (Fig. 2D), suggesting that loss of Oxa1L intensifies MPP^+^-induced mitochondrial pathway disruption.

**Figure 2.**
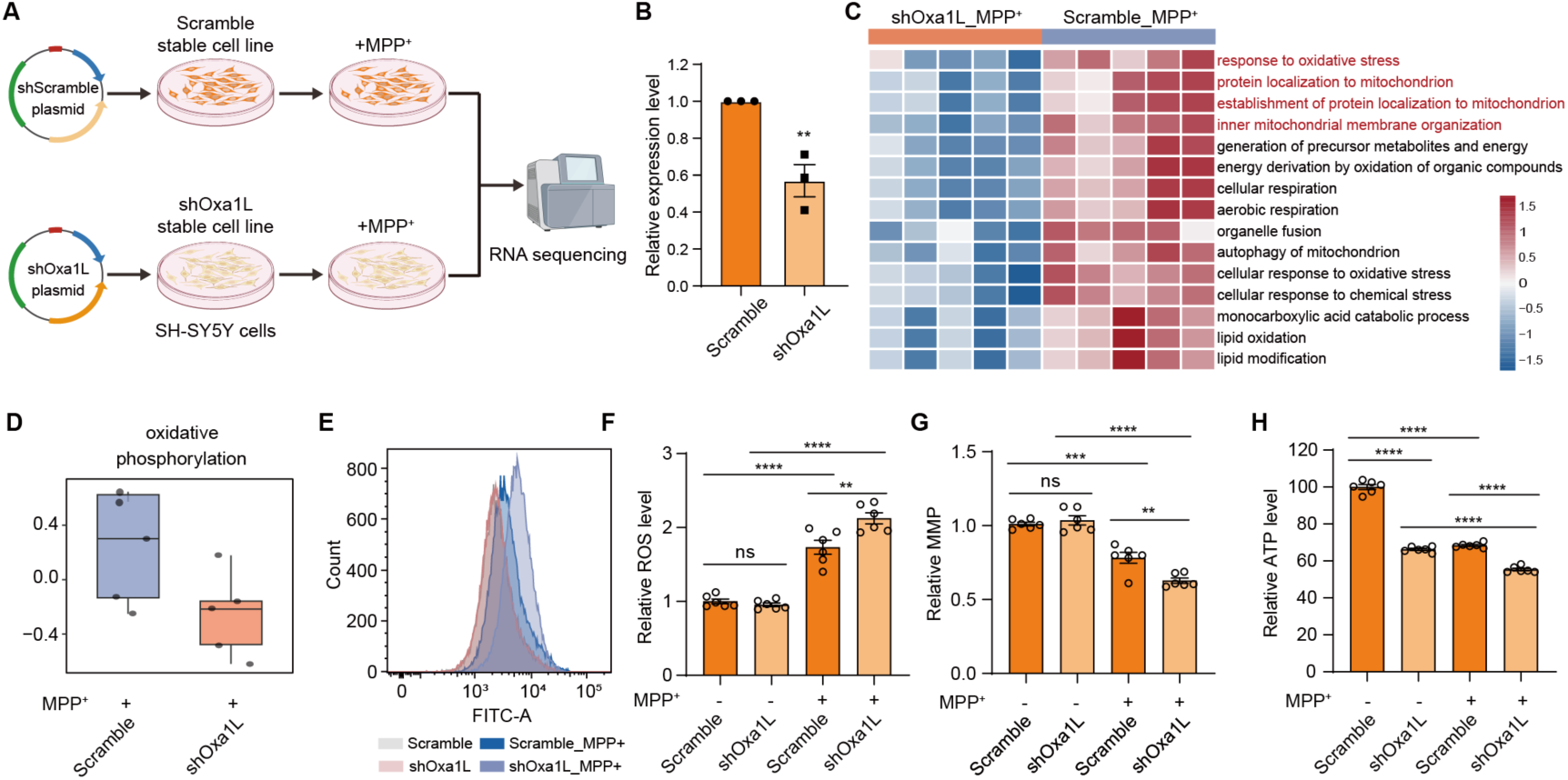
Loss of Oxa1L sensitizes mitochondria to Parkinsonian stress. **A** Schematic of RNA-seq analysis in scramble and shOxa1L SH-SY5Y cells under MPP^+^ treatment. **B** Relative Oxa1L expression in scramble and shOxa1L cells. **C** GO enrichment analysis of downregulated genes in shOxa1L_MPP^+^ and scramble_MPP^+^ cells. **D** KEGG pathway enrichment showing suppression of oxidative phosphorylation upon Oxa1L knockdown. **E** Representative flow cytometry histograms of ROS-associated fluorescence (FITC). **F-H** Quantification of intracellular ROS (F), mitochondrial membrane potential (G), and ATP levels (H). (Data represent mean ± SEM; statistical significance as indicated, *\*\*\*\*p < 0.0001*; *\*\*\*p < 0.001*; *\*\*p < 0.01*; *\*p < 0.05*; ns not significant).

We further evaluated mitochondrial function directly. Upon MPP^+^ treatment, shOxa1L cells accumulated higher levels of intracellular ROS compared with scramble controls (Fig. 2E, F). In parallel, mitochondrial membrane potential was further diminished in Oxa1L-deficient cells (Fig. 2G). Consistent with these changes, ATP production was more severely reduced in shOxa1L cells than in control cells following MPP^+^ exposure (Fig. 2H). Collectively, these results indicate that Oxa1L deficiency aggravates mitochondrial dysfunction induced by Parkinsonian stress, leading to enhanced oxidative imbalance and impaired energy metabolism.

### Effects of Oxa1L overexpression on mitochondrial function under MPP^+^-induced stress

To complement the loss-of-function analyses, we examined whether elevating Oxa1L expression could counteract mitochondrial dysfunction under Parkinsonian stress. A doxycycline (Dox)-inducible Oxa1L expression system was introduced into SH-SY5Y cells. Upon Dox treatment, robust induction of Oxa1L was observed, and immunofluorescence analysis revealed strong colocalization of Oxa1L with mitochondria, as indicated by overlap with MitoTracker staining (Fig. 3A-C).

**Figure 3.**
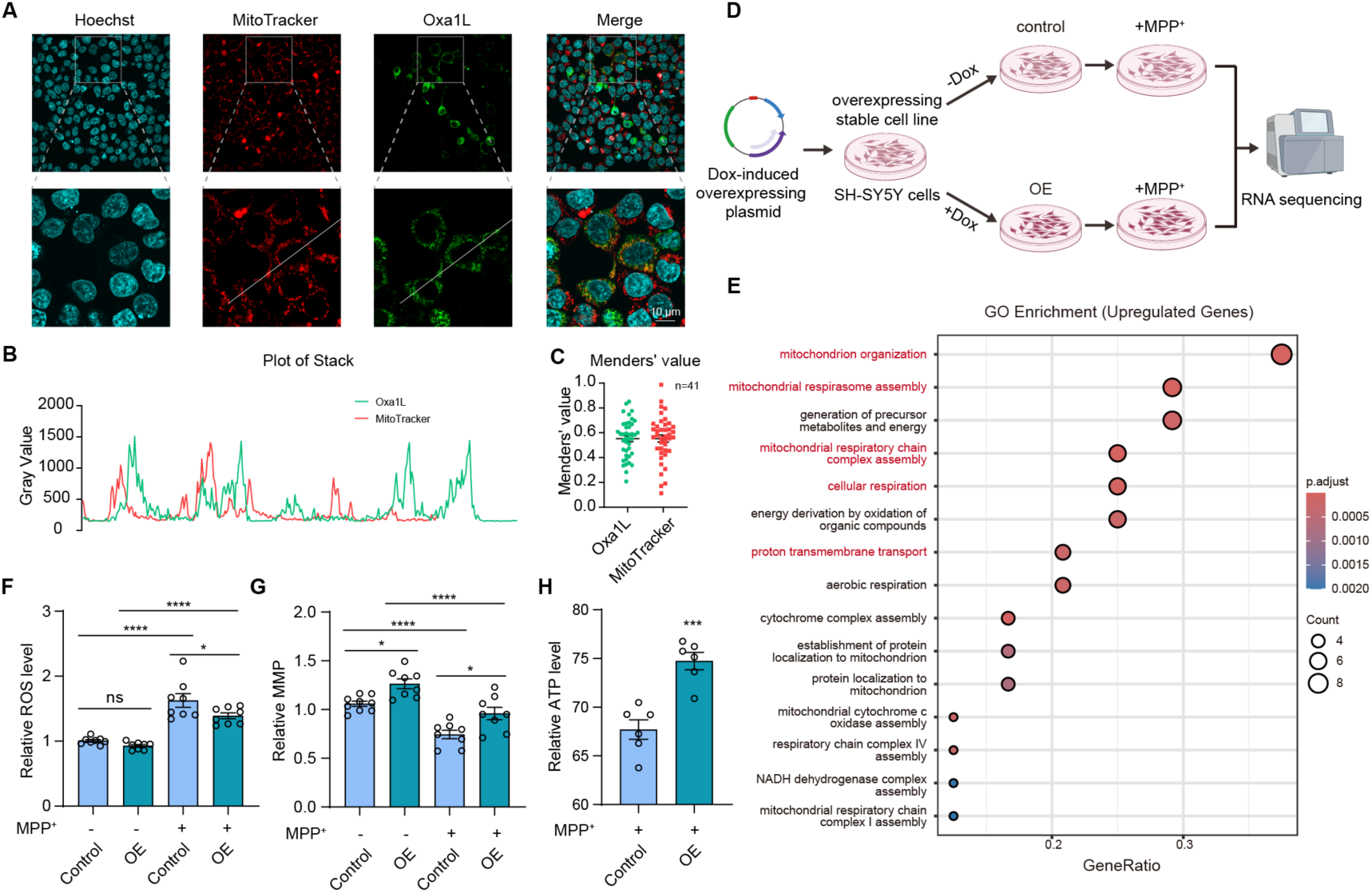
Oxa1L abundance buffers mitochondrial vulnerability under stress. **A.** Immunofluorescence images showing mitochondrial localization of doxycycline-inducible Oxa1L in SH-SY5Y cells. Nuclear was stained with Hoechst, mitochondria with MitoTracker, and Oxa1L with Flag-FITC antibody. **B** Line-scan intensity profiles across the indicated region showing the spatial overlap between Oxa1L and MitoTracker signals. **C** Quantification of colocalization between Oxa1L and mitochondria using Manders’ coefficient. **D** Schematic illustrating the RNA-seq analysis performed in Dox-inducible stable Oxa1L overexpression (OE) cells. **E** GO enrichment analysis of upregulated genes in OE cells compared with control cells, , highlighting pathways related to mitochondrial organization, respiratory chain assembly, and cellular respiration. **F-H** Quantification of intracellular ROS levels (F), mitochondrial membrane potential (G), and ATP levels (H) in control and Oxa1L-overexpressing cells following MPP^+^ treatment. Control, without Dox treatment. OE, treated with 100 ng/ml Dox. (All bar graphs represent mean ± SEM, \*\*\*\**p* < 0.0001; \*\*\**p* < 0.001; \*\**p* < 0.01; \**p* < 0.05; ns not significant).

We subsequently established stable Dox-inducible Oxa1L overexpression (OE) cell lines and performed RNA-seq analysis. GO enrichment analysis revealed the upregulation of pathways related to mitochondrial organization, respiratory chain assembly, cellular respiration, and proton transmembrane transport in Oxa1L-overexpressing cells compared with controls (Fig. 3D, E), suggesting an improved mitochondrial structural and functional capacity.

Functional assays supported these transcriptomic changes. Following MPP^+^ exposure, Oxa1L-overexpressing cells exhibited lower ROS accumulation, preserved mitochondrial membrane potential, and higher intracellular ATP levels relative to control cells (Fig. 3F–H). Together, these findings indicate that increased Oxa1L expression alleviates mitochondrial dysfunction induced by Parkinsonian stress and enhances mitochondrial bioenergetic capacity.

### Selective effects of Oxa1L on mitochondrial genome–encoded OXPHOS subunits

Given the transcriptional and functional effects of Oxa1L manipulation, we next examined mitochondrial protein expression. In wild-type SH-SY5Y cells, MPP^+^ treatment reduced the abundance of several mitochondrially encoded OXPHOS subunits, including MT-CYTB, MT-CO1, MT-CO2, and MT-ATP8 (Fig. 4A). Notably, this reduction was more pronounced in Oxa1L-deficient cells under MPP^+^ exposure (Fig. 4B). In contrast, expression levels of nuclear-encoded respiratory chain subunits, such as NDUFB8, SDHA, and ATP5A1, remained largely unchanged upon Oxa1L knockdown (Fig. 4B). These observations indicate that Oxa1L preferentially influences the expression of mitochondrial genome-encoded proteins, consistent with its role in co-translational insertion at the mitochondrial inner membrane.

**Figure 4.**
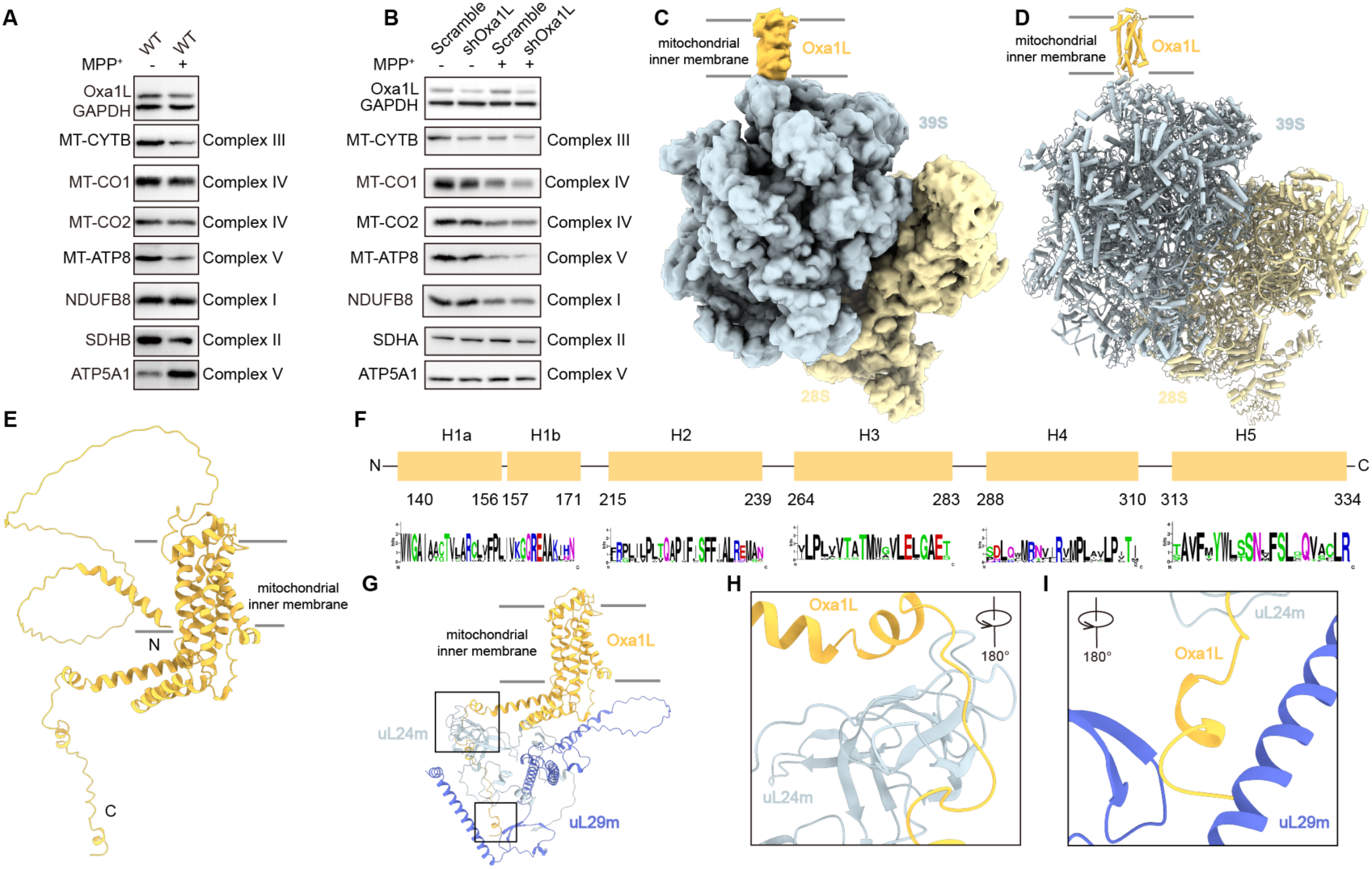
Oxa1L selectively couples mitochondrial translation to OXPHOS assembly. **A** Immunoblot blot analysis of Oxa1L and mitochondrial OXPHOS complex subunits in control and MPP^+^-treated SH-SY5Y cells. Mitochondrial genome-encoded subunits (MT-CYTB, MT-CO1, MT-CO2, MT-ATP8) and nuclear genome-encoded subunits (NDUFB8, SDHB, ATP5A1) are indicated. **B** Immunoblot analysis of OXPHOS subunits in scramble and Oxa1L knockdown (shOxa1L) cells following MPP^+^ treatment, showing selective reduction of mitochondrial genome–encoded proteins upon Oxa1L deficiency. **C-D** Cryo-EM density map (C) and corresponding atomic model (D) of the Oxa1L-mitoribosome complex , revealing Oxa1L positioned adjacent to the ribosomal exit tunnel. **E** AlphaFold3-predicted structure of monomeric human Oxa1L, highlighting its five transmembrane helices. **F** Sequence alignment of Oxa1L transmembrane regions across representative species, illustrating strong evolutionary conservation. Multiple sequence alignment was performed using WebLogo. **G** AF3-predicted model of Oxa1L-uL24m-bL29m complex. The black box indicates the focused area shown in (H, I). **H, I** Enlarged views of interaction interfaces between Oxa1L and ribosomal proteins uL24m (H) and bL29m (I), identifying conserved matrix-exposed regions implicated in co-translational membrane insertion.

To explore the molecular basis of this selectivity, we isolated Oxa1L–mitochondrial ribosome complexes from HEK293F cells and analyzed them by cryo-electron microscopy. Oxa1L was detected as a distinct density positioned adjacent to the ribosomal exit tunnel, near ribosomal proteins mL45 and uL24m (Fig. 4C, D). AlphaFold3-based structural modeling revealed that Oxa1L is a five-pass transmembrane protein with both N- and C-termini oriented toward the mitochondrial matrix (Fig. 4E). The transmembrane regions showed strong evolutionary conservation across species (Fig. 4F). Further structural analysis indicated direct interactions between Oxa1L and ribosomal proteins uL24m and bL29m. Two matrix-exposed regions of Oxa1L, spanning residues 374-406 and 425-435, correspond well with previously reported interaction sites (Fig. 4G–I), suggesting that these regions may mediate coupling between the mitochondrial ribosome and the inner membrane during co-translational insertion ^30^.

### Functional mapping of Oxa1L regions involved in mitochondrial protein biogenesis

To functionally validate the structural insights, we performed a truncation analysis targeting conserved regions of Oxa1L. Seven truncation mutants were generated, including △61-67, △204-213, △212-221, △366-398, △374-406, △373-412, and △425-435, encompassing both predicted transmembrane-proximal and matrix-exposed regions implicated in ribosome interaction. To exclude the possibility that functional defects resulted from impaired mitochondrial targeting, all truncation variants were expressed in SH-SY5Y cells and examined by immunofluorescence microscopy. Each mutant exhibited robust colocalization with the outer mitochondrial membrane marker TOM20, comparable to that observed for full-length Oxa1L (Fig. 5A, B) , indicating that truncation did not disrupt mitochondrial localization.

**Figure 5.**
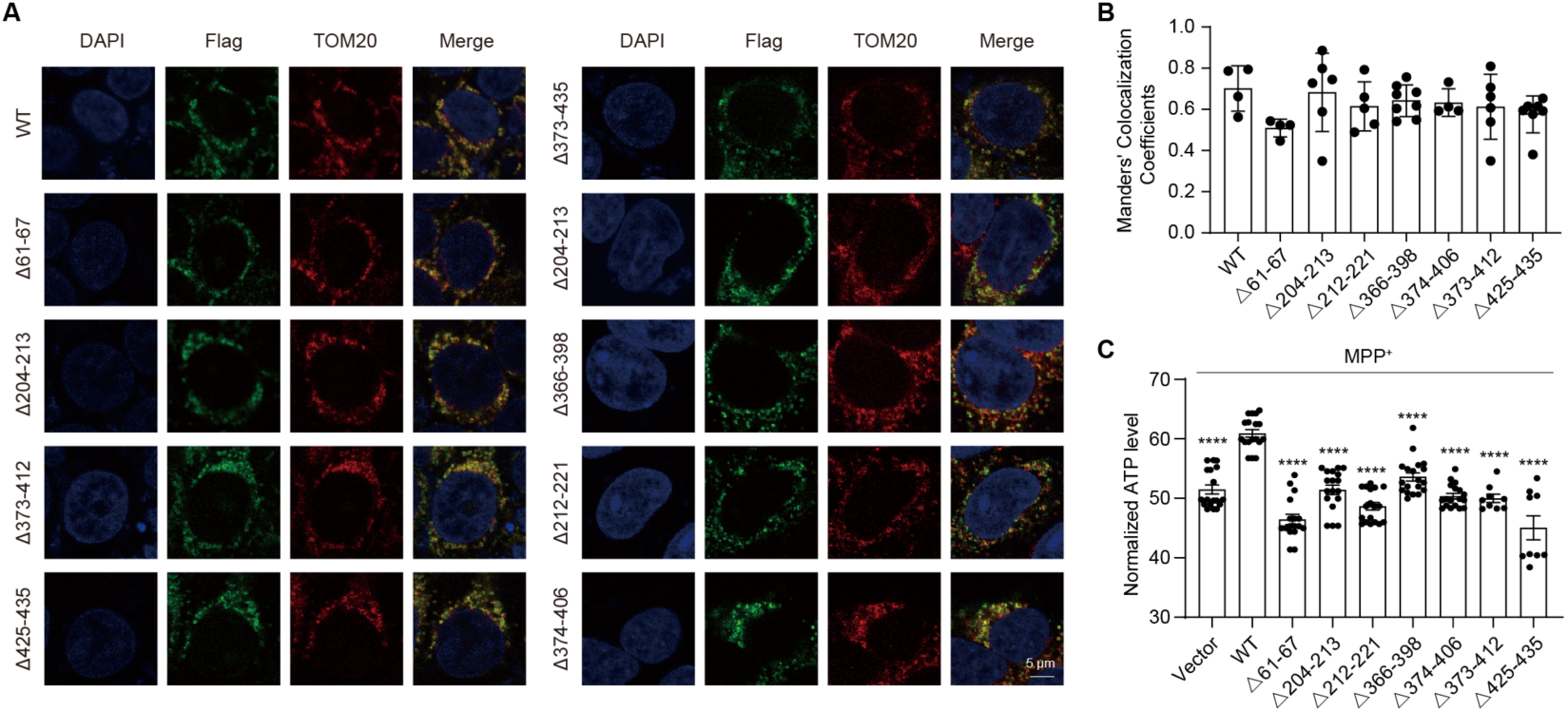
Distributed Oxa1L domains underpin mitochondrial protein insertion. **A** Confocal image showing mitochondrial localization of Flag-tagged Oxa1L truncation mutants in SH-SY5Y cells. Oxa1L truncations were detected by Flag-FITC, mitochondria by TOM20, and nuclei by DAPI. **B** Quantification of Manders’ colocalization coefficients between Oxa1L truncation mutants and TOM20, indicating intact mitochondrial targeting. **C** Cellular ATP levels in cells expressing wild-type (WT) Oxa1L or truncation mutants under MPP^+^ treatment. Loss of ATP maintenance in truncation mutants indicates that multiple Oxa1L regions are involved in mitochondrial protection under stress conditions. (All bar graphs above represent mean ± SEM. *\*\*\*\*p < 0.0001*; *\*\*\*p < 0.001*; *\*\*p < 0.01*; *\*p < 0.05*)

We then evaluated the effect of these Oxa1L truncation mutants to support mitochondrial bioenergetics under Parkinsonian stress. Upon MPP^+^ treatment, cells expressing wild-type Oxa1L maintained higher intracellular ATP levels, whereas none of the truncation mutants were able to preserve ATP production under the same conditions (Fig. 5C). Notably, truncations affecting regions predicted to mediate ribosome interaction failed to confer any detectable bioenergetic protection, despite correct mitochondrial targeting. Together, these analyses identify multiple discrete regions of Oxa1L that contribute to mitochondrial bioenergetic maintenance under stress. Loss of individual structural elements is sufficient to compromise Oxa1L-mediated maintenance of mitochondrial bioenergetics under stress conditions, highlighting the requirement for an intact Oxa1L architecture to support functional coupling between mitochondrial translation and membrane insertion.

## Discussion

Mitochondrial oxidative phosphorylation relies on the precise coordination between the synthesis of mitochondrially encoded proteins and their insertion into the inner membrane. Disruption of this coupling is increasingly recognized as a key contributor to mitochondrial dysfunction under stress, yet the molecular determinants governing this process remain poorly defined. In this study, we identify the mitochondrial insertase Oxa1L as a critical regulator of co-translational membrane insertion that safeguards mitochondrial bioenergetic integrity under Parkinsonian stress conditions.

Our functional analyses reveal that Oxa1L plays an active role in maintaining mitochondrial membrane potential, limiting oxidative stress, and sustaining ATP production during stress. Importantly, both gain- and loss-of-function experiments establish that Oxa1L is not merely a downstream marker of mitochondrial damage, but instead functions as a determinant of mitochondrial vulnerability. These findings position Oxa1L as a modulatory node through which stress can selectively compromise mitochondrial function.

A key insight from our study is the selective sensitivity of mitochondrially encoded respiratory chain subunits to perturbations in Oxa1L function. While nuclear-encoded subunits are largely preserved, the abundance of mitochondrial DNA–encoded proteins is markedly reduced upon Oxa1L impairment. This selectivity suggests that co-translational membrane insertion constitutes a rate-limiting step in mitochondrial protein biogenesis, rendering mitochondrially encoded subunits uniquely vulnerable to reductions in insertion capacity. In this context, Oxa1L emerges as a buffering factor that determines whether mitochondrial translation outputs can be productively incorporated into the inner membrane under stress.

Our structural analyses further provide a spatial framework for understanding Oxa1L function. Localization of Oxa1L at the mitoribosomal exit site supports a model in which translation and membrane insertion are physically coupled, enabling efficient channeling of nascent polypeptides into the inner membrane. Although the current structures do not capture dynamic insertion intermediates, they define an architectural organization that is consistent with Oxa1L acting as an insertion platform rather than a passive membrane anchor.

While our experiments were conducted in Parkinsonian stress models, the mechanisms uncovered here are likely to reflect a general principle governing mitochondrial protein biogenesis under stress. Given the conserved nature of Oxa1-family insertases, limitations in co-translational insertion capacity may represent a broadly applicable vulnerability point across diverse physiological and pathological conditions that challenge mitochondrial homeostasis.

Together, our findings uncover a previously underappreciated layer of mitochondrial quality control operating at the level of co-translational membrane insertion. By defining Oxa1L as a key determinant of mitochondrial resilience, this work provides a conceptual framework linking mitochondrial stress to selective respiratory chain failure and highlights co-translational insertion as a critical checkpoint in maintaining mitochondrial function.

## Methods

### Cell culture and transfection

SH-SY5Y (SCSP-5014) cells were obtained from the Cell Bank of the Chinese Academy of Sciences and authenticated by the supplier using STR profiling. Cells were cultured in Dulbecco’s modified Eagle medium (DMEM; Thermo Fisher Scientific) supplemented with 10% (v/v) fetal bovine serum (FBS; Moregate), 100 units/mL penicillin, and 0.1 mg/mL streptomycin (MeilunBio) at 37 °C in 5% CO₂. Stable Oxa1L over-expressed and knockdown SH-SY5Y cells were established by our lab. The human *Oxa1L* gene was cloned into the pLVX-Tet3G vector (Takara, 631847). SH-SY5Y cells were transduced with lentivirus and OXA1L expression was induced with doxycycline (100 ng/mL, 48 h). For knockdown, SH-SY5Y cells were transduced with pLKO.1-based lentivirus (Addgene, 8453) encoding an shRNA targeting human OXA1L (5′-GCAGGAGACCATATTGAGTAT-3′). For selection, puromycin (1 μg/mL) was included in the culture medium. Transient transfections of SH-SY5Y cells were performed with a liposomal transfection reagent (MeilunBio) according to the manufacturer’s instructions. Cells were treated with MPP^+^ for 24 hours to induce a PD model.

### Immunoblotting

Proteins were separated by SDS–PAGE (12% gels) and transferred to PVDF membranes (Cytiva). Membranes were probed with primary antibodies against DYKDDDDK (FLAG) (Flag tag, proteintech, 1:5000), GAPDH (proteintech, 1:5000), Oxa1L (proteintech, 1:5000), MT-CYTB (proteintech, 1:2000), MT-CO1 (Cell Signaling Technology, 1:1000), MT-CO2 (proteintech, 1:2000), MT-ATP8 (abclonal, 1:1000), NDUFB8 (Cell Signaling Technology, 1:1000), SDHA (abclonal, 1:1000), SDHB (Cell Signaling Technology, 1:1000), and ATP5A1 (abclonal, 1:10000). The second antibody is Peroxidase AffiniPure Goat Anti-Mouse IgG (H + L) (YEASEN, 1:10000) and Peroxidase AffiniPure Goat Anti-Rabbit IgG (H + L) (YEASEN, 1:10000). The relative protein expression levels were calculated using Fiji software.

### qRT-PCR

Total RNA was isolated from SH-SY5Y cells using an RNA isolation kit (Vazyme). First-strand cDNA was synthesized with the RT SuperMix kit (Vazyme), which includes a genomic DNA removal step. Quantitative real-time PCR was performed on a StepOnePlus Real-Time PCR System (Thermo Fisher Scientific) using SYBR probes.

### Immunofluorescence and confocal imaging

SH-SY5Y cells were cultured on poly-D-lysine coated glass coverslips for 24 h prior to transfection. Subsequently, the cells were transfected with Flag-tagged Oxa1L plasmids (WT or mutants) for 16 h. Samples were then fixed using 4% PFA, permeabilized with Triton X-100, and blocked in a solution containing 2% BSA in PBST. Primary antibody incubation was carried out overnight at 4 °C using a mouse anti-Flag antibody (proteintech, 1:5000) and a rabbit anti-TOM20 antibody (proteintech, 1:200). After washing three times with PBS, the samples were probed with a FITC-conjugated anti-mouse secondary antibody (Beyotime, 1:200) and a Cy3-conjugated anti-rabbit secondary antibody (Beyotime, 1:200) for 1 h at room temperature. Nuclei were stained with DAPI (Yeasen). Finally, coverslips were mounted using an antifade medium. Confocal imaging was performed on an Olympus FV3000 system equipped with a 60× oil immersion objective, capturing representative images from at least three random fields.

### Mitochondrial membrane potential measurement

Mitochondrial membrane potential was assessed using the JC-1 probe. SH-SY5Y cells were exposed to 1 mM MPP^+^ for 24 h, harvested, and incubated with JC-1 at 37 °C for 30 min. Cells were then washed twice with cold JC-1 buffer and analyzed on a flow cytometer (FACS Fortessa, BD) using FITC and PE channels. Fluorescence data were processed in FlowJo v10.8.1.

### ROS detection

Intracellular reactive oxygen species (ROS) were quantified using the DCFH-DA probe. SH-SY5Y cells were treated with 1 mM MPP^+^ for 24 h, harvested, and incubated with 10 μM DCFH-DA at 37 °C for 30 min. Cells were washed twice with PBS and analyzed by flow cytometry (FACS Fortessa, BD). Data were processed in FlowJo v10.8.1.

### ATP detection

Cellular ATP was quantified using the CellTiter-Meiluncell Luminescent Reagent (MeilunBio) following the manufacturer’s instructions. SH-SY5Y cells were seeded in 96-well plates and treated with 1 mM MPP^+^ for 24 h. Plates were equilibrated to room temperature for 10 min, after which the luminescent reagent was added and incubated for an additional 10 min at room temperature. Luminescence was recorded on a Spark microplate reader (Tecan), and relative ATP levels were calculated. Data analysis was performed in GraphPad Prism.

### Isolation of mitochondria

Cells were harvested at 4 °C, washed once with cold PBS, and resuspended in mitochondrial isolation buffer (MIB; 50 mM HEPES-KOH pH 7.4, 10 mM KCl, 1.5 mM MgCl₂, 1 mM EDTA, 1 mM EGTA, 1 mM DTT, protease inhibitors). After swelling for 15 min with gentle stirring at 4 °C, one-third volume of SM4 buffer (280 mM sucrose, 840 mM mannitol, 50 mM HEPES-KOH pH 7.5, 10 mM KCl, 1.5 mM MgCl₂, 1 mM EDTA, 1 mM EGTA, 1 mM DTT, protease inhibitors) was added, and cells were disrupted in a pre-chilled Teflon/glass Dounce homogenizer (>100 strokes). The homogenate was cleared at 800 × g, 15 min, 4 °C, and the supernatant was passed through Miracloth. The pellet was re-homogenized in MIB:SM4 (3:1), centrifuged again at 800 × g, and the combined supernatants were clarified at 1,000 × g (15 min, 4 °C) followed by 10,000 × g (15 min, 4 °C) to pellet crude mitochondria. The pellet was gently rinsed, resuspended in MIB:SM4 (3:1), supplemented with RNase-free DNase I (200 U per 10 g wet cell pellet), and rotated for 20 min at 4 °C. After centrifugation at 10,000 × g (15 min, 4 °C), mitochondria were resuspended in SEM buffer (250 mM sucrose, 1 mM EDTA, 20 mM MOPS-KOH pH 7.4), gently Dounced (≤5 strokes), and layered onto a discontinuous sucrose step gradient (15/23/32/60% in 20 mM HEPES-KOH pH 7.4, 1 mM EDTA). Gradients were spun in an SW40 rotor at ∼28,000 × g for 1 h at 4 °C. Intact mitochondria were collected from the brown band at the 32%/60% interface, transferred to a clean tube, and either used immediately or snap-frozen in liquid nitrogen and stored at −80 °C.

### Oxa1L-mitoribosome complex extraction

Oxa1L-mitoribosome complex was extraction from mitochondrial as described in our previous studies^19, 35^. Frozen mitochondria were thawed on ice and lysed by adding two volumes of detergent lysis buffer (25 mM HEPES-KOH pH 7.4, 150 mM KCl, 50 mM Mg(OAc)₂, 1.5% n-dodecyl-β-D-maltoside, 0.15 mg/mL cardiolipin, 250–500 mM GMPPCP, 2 mM DTT, protease inhibitors). Samples were mixed by inversion, briefly Dounced to assist solubilization, and rotated 20 min at 4 °C. Insoluble material was removed by centrifugation at ∼30,000 × g for 20 min at 4 °C. The supernatant was carefully layered over a 1 M sucrose cushion (20 mM HEPES-KOH pH 7.4, 100 mM KCl, 20 mM Mg(OAc)₂, 0.6% n-dodecyl-β-D-maltoside, 0.06 mg/mL cardiolipin, 250 mM GMPPCP, 2 mM DTT) and centrifuged at ∼231,550× g for 60 min at 4 °C. Pellets were gently rinsed with resuspension buffer (20 mM HEPES-KOH pH 7.4, 100 mM KCl, 5 mM Mg(OAc)₂, 0.15% n-dodecyl-β-D-maltoside, 0.015 mg/mL cardiolipin, 250 mM GMPPCP, 2 mM DTT) and resuspended in ∼100 µL of the same buffer. The absorbance at 260 nm was recorded, and the entire sample was applied to a 15–30% linear sucrose gradient (in 20 mM HEPES-KOH pH 7.4, 100 mM KCl, 5 mM Mg(OAc)₂, 0.05% n-dodecyl-β-D-maltoside, 0.005 mg/mL cardiolipin, 250 mM GMPPCP, 2 mM DTT). Gradients were centrifuged in a TLS-55 rotor at ∼213,626 × g for 60–90 min at 4 °C, fractionated, and nucleic-acid–containing fractions (A₂₆₀/A₂₈₀ > 1.6) were pooled. The sample was concentrated and the absorbance at 260 nm (A260) was measured until it reached 4-6.

### Cryo-EM sample preparation and data acquisition

Aliquots of 3.3 μl ∼4-6 mg/mL extracted Oxa1L-mitoribosome samples were applied to glow-discharged holey carbon grids (Quantifoil, R2/2, Cu+2nm C, 300 mesh). The grids were blotted for 4.5-5.5 s using Vitrobot at 4°C with 100% humidity, and plunge-frozen into the liquid ethane.

Cryo-EM datasets were collected on Titan Krios G4 cryo-electron microscope operated at 300 kV, equipped with a Falcon 4i Direct Electron Detector and a Selectris X energy filter (Thermo Fisher Scientific). Movie stacks were automatically collected using EPU at a magnification of 130,000× with a pixel size of 0.932 Å for a total dose per EER (electron event representation) movie of ∼50 e^−^/Å^2^. The defocus range was set between -0.8 to -1.8 µm.

### Data processing

All datasets were processed in CyroSPARC (v4.5.3)^36^. All of the movies were initially aligned using Patch Motion Correction and contrast transfer function (CTF) was determined using patch CTF determination. Particles were auto-picked using the blob picker and template picker. Particles were extracted and subjected to one round of two-dimensional (2D) classification to remove junk particles. Good classes were then used for ab initio reconstruction. Particles from classes displaying membrane protein characteristics and their corresponding cryo-EM maps were refined using non-uniform refinement, resulting in a consensus map. Multiple rounds of heterogeneous refinement and three-dimensional (3D) classification were performed. Selected particles were subjected to non-uniform (NU) refinement and local refinement with C1 symmetry imposed with a mask covering the Oxa1L blob. The models of mitoribosome were automated refined in PHENIX by using Phenix.real_space_refine^37^. Model of Oxa1L was generated by AlphaFold3^29^. The predicted Oxa1L model was fitted into the mitoribosome using UCSF ChimeraX^38^.

## Acknowledgments

We thank Yuli Jiang and Yi Lv from the platform of the Institute for Translational Brain Research at Fudan University for their support with imaging, molecular, and animal experiments. We thank Zhenguo Chen, Anqi Dong, and Hui Zhao at the Center of cryo-EM at Fudan University for their support with cryo-EM data collection. This work was supported by the grants from the National Natural Science Foundation of China (32171216 to HY), China Postdoctoral Science Foundation (2023M730689 to XS), Fudan Undergraduate Research Opportunities Program (23216 to RH), and the Zhengyi Scholar Research Concluding Project from the School of Basic Medical Sciences (Fudan University, S32-B03 to YZ).

## Author Contributions

Conceptualization: HY, XS

Methodology: XS, RH, HZ

Investigation: XS (flow cytometry, ATP detection, RNAseq sample preparation and analysis, confocal imaging and analysis, stable cell line optimization, Oxa1L-ribosome complex extraction, cryo-EM sample preparation, data collection and processing)

RH (flow cytometry, ATP detection, RNAseq analysis, stable cell line optimization, Oxa1L-ribosome complex extraction, Western blot, truncation mutagenesis construction)

HZ (stable cell line optimization, Western blot, confocal imaging, and analysis) PS (structure analysis)

JKZ (cryo-EM data processing)

YZ (ATP detection, truncation mutagenesis construction)

Visualization: XS, RH, HZ, PS

Supervision: HY

Writing—original draft: XS, RH, HY

Writing—review & editing: XS, HY

